# Divergence in Population Coding for Space between Dorsal and Ventral CA1

**DOI:** 10.1101/2021.03.10.434839

**Authors:** Udaysankar Chockanathan, Krishnan Padmanabhan

**Affiliations:** Neuroscience Graduate Program, University of Rochester School of Medicine and Dentistry, Rochester, NY 14623, USA; Medical Scientist Training Program, University of Rochester School of Medicine and Dentistry, Rochester, NY 14623, USA; Ernest J. Del Monte Institute for Neuroscience, University of Rochester School of Medicine and Dentistry, Rochester, NY 14623, USA; Department of Neuroscience, University of Rochester School of Medicine and Dentistry, Rochester, NY 14623, USA; Center for Visual Science, University of Rochester School of Medicine and Dentistry, Rochester, NY 14623, USA

**Author notes:** Correspondence (K.P.).

## Abstract

Molecular, anatomic, and behavioral studies show that the hippocampus is structurally and functionally heterogeneous, with dorsal hippocampus implicated in mnemonic processes and spatial navigation and ventral hippocampus involved in affective processes. By performing electrophysiological recordings of large neuronal populations in dorsal and ventral CA1 in head-fixed mice navigating a virtual environment, we found that this diversity resulted in different strategies for population coding of space. We found that the populations of neurons in dorsal CA1 had a higher dimensionality and showed more complex patterns of activity, translating to more information being encoded, as compared ensembles in vCA1. Furthermore, a pairwise maximum entropy model was better at predicting the structure of global patterns of activity in ventral CA1 as compared to dorsal CA1. Taken together, we uncovered different coding strategies that likely emerge from anatomical and physiological differences along the longitudinal axis of hippocampus and that may, in turn, underpin the divergent ethological roles of dorsal and ventral CA1.

**Highlights:**

- No differences in dCA1 and vCA1 place field size in recordings of neuronal populations in mice navigating a virtual environment
- dCA1 has higher single-neuron and population-level spatial information compared to vCA1 due to differences in the sparsity of firing.
- Population activity of dCA1 has higher entropy and is higher dimensional than vCA1
- Pairwise maximum entropy models are better at predicting population activity in vCA1 compared to dCA1

## Introduction

The hippocampus is a functionally diverse brain region, linked to an array of cognitive and emotional behaviors (Klüver and Bucy, 1937; Scoville and Milner, 1957). This diversity is particularly pronounced along the dorsal-ventral axis of CA1, with dorsal CA1 implicated in episodic memory and spatial navigation in contrast to ventral CA1, which has been linked to anxiety and social cognition (Fanselow and Dong, 2010; Henke, 1990; Kjelstrup et al., 2002; Moser and Moser, 1998; Moser et al., 1995; Okuyama et al., 2016).

Functional diversity arises from differences at multiple scales in CA1, including differences in gene expression (Cembrowski et al., 2016; Thompson et al., 2008) to variations in intrinsic neuronal biophysical and morphologic properties (Dougherty et al., 2012, 2013; Malik et al., 2016) to diverse afferent and efferent anatomical connections (Cenquizca and Swanson, 2007; Meira et al., 2018; Padmanabhan et al., 2019; Swanson and Cowan, 1977). While these intrinsic and circuit-level differences contribute to the array of diverse behaviors linked to dorsal and ventral hippocampus, they also underlie differences of a single representation along the dorsal-ventral axis of the hippocampus. For example, place cells, neurons that preferentially fire when an animal visits a particular region of its environment (O’Keefe and Dostrovsky, 1971), are present throughout the dorsal-ventral axis of CA1. The place fields became progressive larger along this axis, with neurons in ventral CA1 providing less information about the animal’s position than those in dorsal CA1 (Ciocchi et al., 2015; Jung et al., 1994). As place cells are considered a critical component of the cognitive map, this suggests that the internal representation of space coarsens in ventral CA1 relative to dorsal CA1. However, multiple studies in dorsal hippocampus have demonstrated that the fidelity of spatial coding is not a simple function of place field size.

First, not all CA1 neurons are place cells. Numerous spatial navigation studies have demonstrated that the pool of place cells varies between environments, but that in any given environment, a subset of dorsal CA1 neurons do not show a place field (Rich et al., 2014; Wilson and McNaughton, 1993). Moreover, while most studies of place cells define a threshold spatial information or sparsity (Skaggs et al., 1993), there is not a sharp boundary between place and non-place cells; dorsal CA1 neurons have variable amounts of spatial information and an animal’s position in space is likely encoded in combination with other variables, such as time, and reward (Gauthier and Tank, 2018; Haimerl et al., 2019; Stefanini et al., 2020). Activity of place cells is determined not only by the position of the animal, but also by correlations with other neurons, including non-place cells; these contributions from the broader population were able to partially explain the trial-to-trial variability in place cell responses (Meshulam et al., 2017). Neurons without clear place fields can thus be highly valuable for spatial coding, and place cells with high spatial information are not necessarily the most useful neurons in the population for decoding position (Stefanini et al., 2020).

Despite accumulating evidence that position information in the hippocampus is collectively encoded by groups of neurons, relatively little is known about how this population activity varies across the dorsal-ventral axis of CA1. Does the activity of the overall population compensate for this larger single-cell place fields in a way that maintains the precision of spatial representations across dorsal and ventral CA1? Alternatively, does the resolution of spatial coding in populations of neurons become coarser in ventral CA1, as it does for the place fields of individual neurons, suggesting divergent computational strategies for encoding place?

By recording from large neuronal populations from both regions as animals navigated a virtual track, we found a divergence in the population representation of space between dorsal and ventral CA1. Both at the single-neuron level and the population level, activity in dorsal CA1 was more informative about the animal’s position than that in ventral CA1. This increased spatial information was underpinned by more complex patterns of population activity in dorsal CA1 that increased the dimensionality of the neural code. To understand how these complex activity patterns may arise, we fit maximum entropy models to the data, which revealed that population-wide patterns could be predicted better using pairwise interactions between neurons in ventral CA1 as compared to dorsal CA1. Taken together, these results suggest that differences in the functional interactions between neurons across the longitudinal hippocampal axis result in differential coding strategies and divergent representations of space in dorsal and ventral CA1.

## Results

### Head-fixed Dorsal and Ventral CA1 Population Recordings in Virtual Reality

To compare the features of neuronal population activity across the dorsal-ventral axis of CA1 hippocampus, we performed electrophysiological extracellular recordings in awake head-fixed mice on a running wheel in a virtual environment. High-density 128-channel silicon electrode arrays were targeted to either dorsal CA1 or ventral CA1 of four male C57BL6/J mice of 9-10 weeks of age while they ran through a virtual one-dimensional virtual track (Figures 1A and 1B; see Method Details) (Gauthier and Tank, 2018). Recording sessions lasted approximately one hour, during which the mice completed multiple runs along the track (mean±std: 37±22 laps per recording session; Figures 1C and 1D). All locomotion was self-motivated; the wheel was non-motorized and no reward was provided.

**Figure 1.**
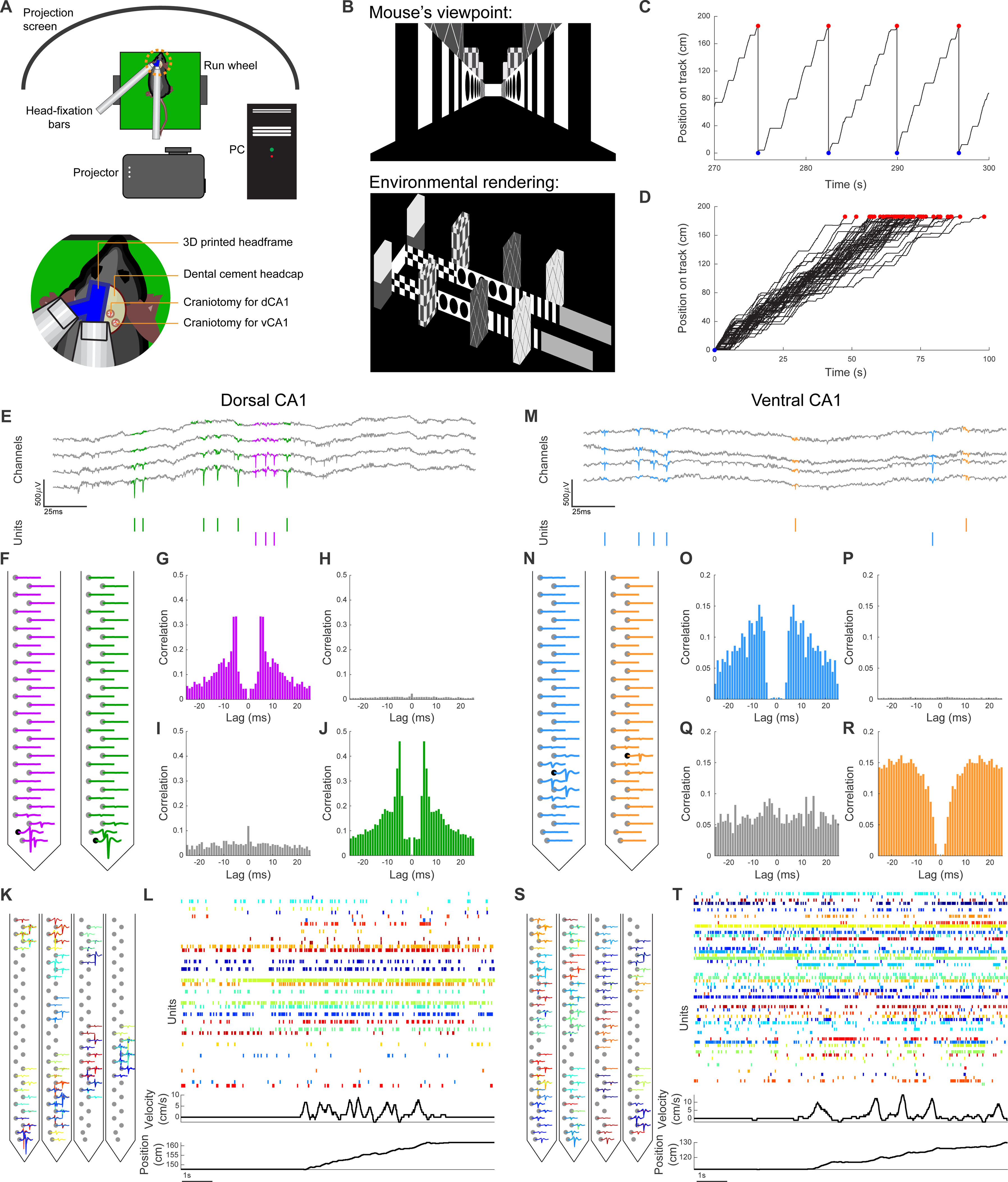
Head-fixed Recordings of Neuronal Populations in Virtual Reality. (A) Schematic of recording setup with virtual environment. (B) Top: Sample frame of virtual environment, shown from the perspective of the mouse. Bottom: Schematic of 1.9m virtual track. (C) Sample of animal trajectory through virtual track. When mice reached end of track (red circles), they were transported to the start of the track (blue circles). (D) Superimposed trajectories of all laps completed in one representative recording session. (E and M) Top: Sample trace of wide-band (0.1Hz-3500Hz) electrophysiological recording in dorsal CA1 (E) and ventral CA1 (M). Spike waveforms from different units are highlighted in different colors. Bottom: Raster plot denoting spike times for each unit. (F and N) Mean waveform of each unit in (E and M) across all 32 channels on a single shank of the electrode array. (G, J, O, and R) Autocorrelograms for units in (E and M), showing clear refractory periods. (H, I, P, and Q) Cross-correlograms for units in (E and M), showing no refractory period. (K and S) Mean waveform of all units from a single dorsal CA1 (K) or ventral CA1 (S) recording session with each color denoting a single unit. For each unit, waveforms are shown on the channel with the largest amplitude spike as well as three neighboring channels. (L and T) Top: Sample of dorsal CA1 (L) and ventral CA1 (T) population activity. The color of the raster plot row corresponds to the unit waveforms in (K and S). Middle: Simultaneous running velocity of animal. Bottom: Simultaneous position of animal on virtual track.

For both dorsal and ventral CA1, individual action potentials, or spikes, were observed in the broadband voltage traces (Figures 1E and 1M). Template matching with Kilosort (Pachitariu et al., 2016) was used to cluster the spike waveforms and identify putative units across multiple neighboring channels. The resulting units were then curated using Phy (Rossant et al., 2016); only those with asymmetric, spike-like waveforms (Figures 1F and 1N) as well as a clear refractory period in their autocorrelograms (Figures 1G, 1J, 1O, and 1R) were included in analysis. Units were merged based on the similarity of their waveforms and the features of their cross-correlograms (Figures 1H, 1I, 1P, and 1Q) were merged. This process yielded between 6 and 80 units per recording in dorsal CA1 and between 43 and 62 units per recording in ventral CA1 (Figures 1K and 1S). The resulting neuronal populations revealed complex patterns of activity in both regions as the animals navigated the virtual space (Figures 1L and 1T).

### Coarsening of Single-neuron Spatial Representations along Dorsal-Ventral Axis of CA1

Previous studies using rodents in an open field have shown that the neuronal representation of space varies along the dorsal-ventral axis of the hippocampus, with ventral CA1 neurons showing more diffuse place fields and lower spatial information than dorsal CA1 neurons (Ciocchi et al., 2015; Jung et al., 1994; Keinath et al., 2014). To test whether this held true in a virtual environment with head-fixed animals, we first visualized the activity of each neuron as a function of the animal’s position along the virtual track (Figures 2A and 2D). In both dorsal and ventral CA1 neurons, we observed place fields that tiled the length of the track (Figures 2B and 2E).

**Figure 2.**
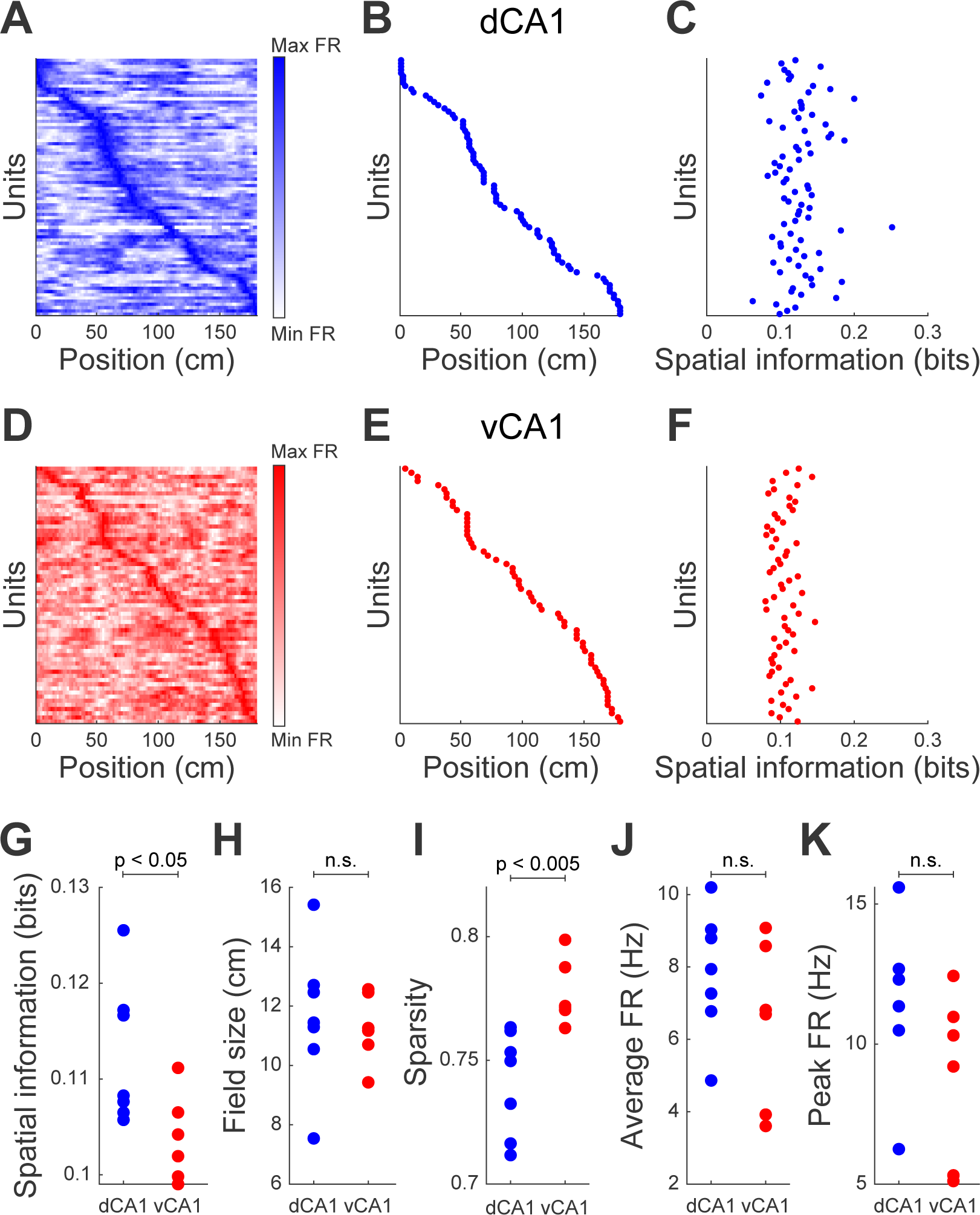
Ventral CA1 Neurons Have Lower Spatial Information than Dorsal CA1 Neurons. (A and D) Map of single-neuron activity as a function of animal position in virtual track for a representative recording session in dorsal CA1 (A) and ventral CA1 (D). Each row denotes a single unit. (B and E) Location in virtual track of peak firing rate for each unit in (A and D). (C and F) Spatial information of each unit in (A and D). (G) Dorsal CA1 neurons had larger spatial information than ventral CA1 neurons (mean ± std: dCA1 = 0.113 ± 0.007 bits, vCA1 = 0.104 ± 0.005 bits, *p* = 0.04, two-sided Wilcoxon rank-sum test, ndCA1 = 7 recording sessions, nvCA1 = 6 recording sessions). Each point denotes a recording session. (H) Place field sizes were not significantly different for dorsal and ventral CA1 neurons (mean ± std: dCA1 = 11.6 ± 2.4 cm, vCA1 = 11.3 ± 1.2 cm, *p* = 0.53, two-sided Wilcoxon rank-sum test, ndCA1 = 7 recording sessions, nvCA1 = 6 recording sessions). Each point denotes a recording session. (I) Dorsal CA1 neurons had sparser activity than ventral CA1 neurons (mean ± std: dCA1 = 0.74 ± 0.02, vCA1 = 0.78 ± 0.01, *p* = 0.002, two-sided Wilcoxon rank-sum test, ndCA1 = 7 recording sessions, nvCA1 = 6 recording sessions). Each point denotes a recording session. (J) Average firing rates were not significantly different for dorsal and ventral CA1 neurons (mean ± std: dCA1 = 7.84 ± 1.74 Hz, vCA1 = 6.45 ± 2.29 Hz, *p* = 0.30, two-sided Wilcoxon rank-sum test, ndCA1 = 7 recording sessions, nvCA1 = 6 recording sessions). Each point denotes a recording session. (K) Peak firing rates were not significantly different for dorsal and ventral CA1 neurons (mean ± std: dCA1 = 11.57 ± 2.83 Hz, vCA1 = 8.89 ± 3.04 Hz, *p* = 0.10, two-sided Wilcoxon rank-sum test, ndCA1 = 7 recording sessions, nvCA1 = 6 recording sessions). Each point denotes a recording session.

We quantified these activity patterns by calculating the spatial information (Figures 2C and 2F) (Skaggs et al., 1993), a measure of how the neuron’s firing rate predicted the animal’s location. Ventral CA1 neurons had a lower information content than dorsal CA1 neurons (Figure 2G), consistent with previous studies on freely behaving animals in open field environments (Jung et al., 1994). Surprisingly, we found that the decreased information content in ventral CA1 neurons was not due to larger place fields, as the place field size was not significantly different between the two regions (Figure 2H). Instead this reduction in spatial information in ventral CA1 was due to a higher level of spontaneous firing at track positions not associated with the place field. This was quantified by the sparsity metric (Skaggs et al., 1996), which showed that ventral CA1 neurons were active over a larger fraction of the track than dorsal CA1 neurons (Figure 2I). Importantly, the difference in spatial information between the two regions could not be explained by differences in either the average firing rates or the peak firing rates, which were not significantly different between ventral and dorsal CA1 neurons (Figures 2J and 2K). Additionally, the relationship between mean firing rates and spatial information was not significantly different for the two regions (Figure S1). Taken together, these results suggest a coarsening of spatial representations from dorsal to ventral CA1 at the level of single neurons in virtual environments was not due to differences in place-field size, but rather to differences in the activity of neurons when the animal was outside of their place fields. Consequently, the reduction of spatial information across the dorsal-ventral axis of CA1 appeared to be a universality principle of neuronal coding, the properties by which this coarsening of information occurred was different in real and virtual environments.

### Preserved Pairwise Correlations, but Weakened Functional Network Structure in Ventral CA1

The differences in single-neuron spatial encoding shown in Figure 2 are consistent with previous literature showing the heterogeneity of neuronal coding properties across the dorsal-ventral axis of CA1 (Ciocchi et al., 2015; Jimenez et al., 2018; Jung et al., 1994; Okuyama et al., 2016). However, these single-cell responses are embedded in a much larger population-level response; these functional interactions between individual neurons shape the collective activity of the CA1 population in ways that are critical for the encoding of position (Meshulam et al., 2017). As our data suggest that place field size alone in virtual environments did not vary, we wished to test whether these functional interactions across neurons in dorsal and ventral CA1 were different (Figures 3A and 3B).

**Figure 3.**
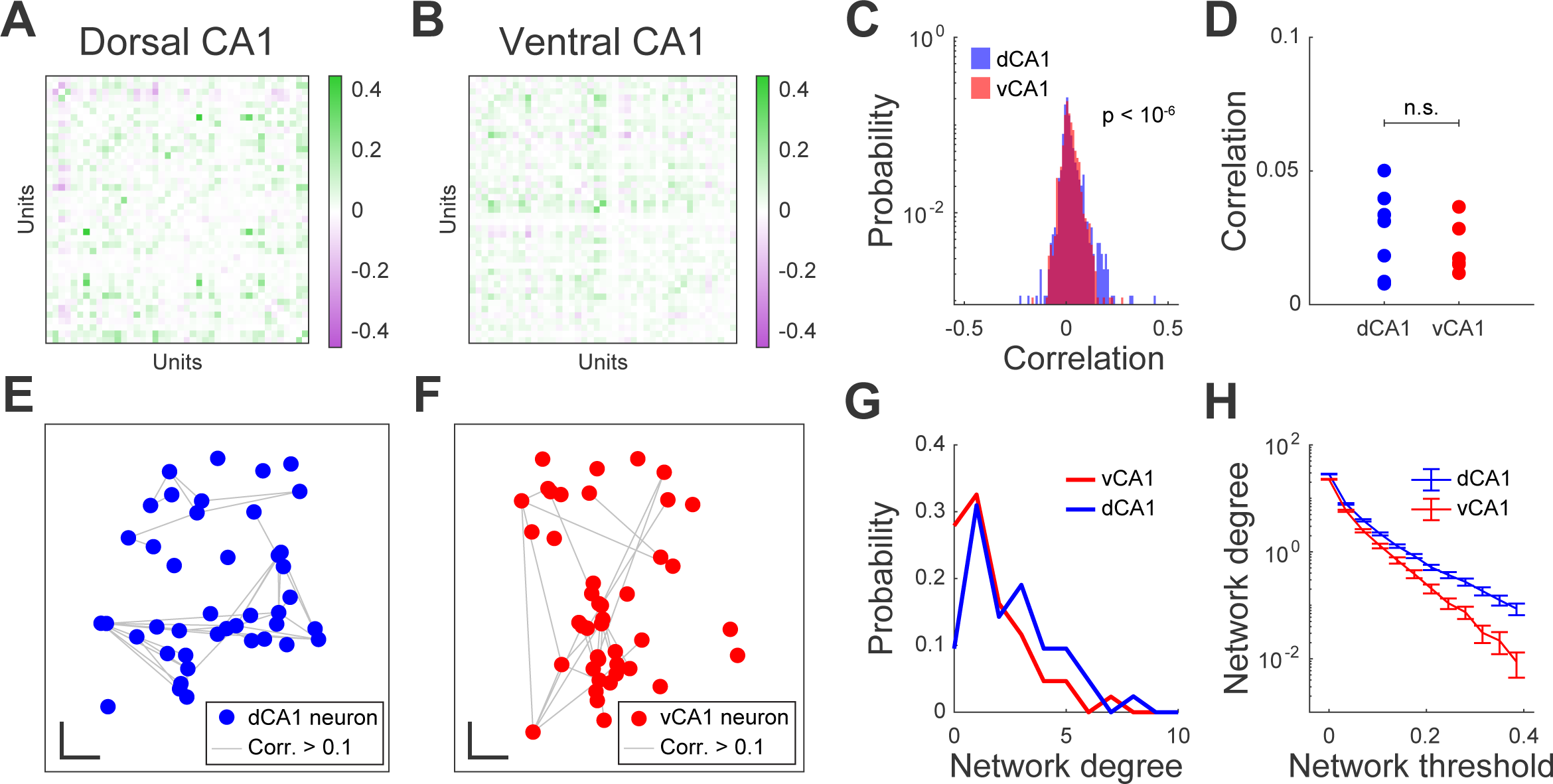
Preserved Correlations, but Weaker Network Structure in Ventral CA1. (A and B) Matrix of Pearson correlation coefficients for every pair of units in a representative recording session in dorsal CA1 (A) and ventral CA1 (B). Each grid cell denotes the correlation coefficient for a single pair of units. (C) Distribution of correlation coefficients for the two example sessions in (A and B). There was no significant difference in the mean correlation between dorsal and ventral CA1 (mean ± std: dCA1 = 0.018 ± 0.049, vCA1 = 0.018 ± 0.038, *p* = 0.06, two-sided Wilcoxon rank-sum test, ndCA1 = 861 pairs of units, nvCA1 = 903 pairs of units). However, the variance of the correlation distribution was larger in dorsal CA1 than in ventral CA1 (variance: dCA1 = 2.4×10^-3^; vCA1 = 1.4×10^-3^, *p* < 10^-6^, two-sided F-test for variance, ndCA1 = 861 pairs of units, nvCA1 = 903 pairs of units). (D) Average correlation coefficients were not significantly different for dorsal and ventral CA1 neurons (mean ± std: dCA1 = 0.027 ± 0.016, vCA1 = 0.021 ± 0.010, *p* = 0.53, two-sided Wilcoxon rank-sum test, ndCA1 = 7 recording sessions, nvCA1 = 6 recording sessions). Each point denotes a recording session. (E and F) Network representation of correlations for a representative dorsal CA1 (E) and ventral CA1 (F) recording session. Points denote the approximate physical position of individual neurons (jitter added to minimize overlapping neurons) and grey lines denote suprathreshold correlations between pairs of neurons. Scale bars denote 100µm in horizontal and vertical directions. (G) Distribution of network degree (number of incident edges) for the representative example networks in (E and F). (H) Mean degree of networks in dorsal CA1 was higher than that of ventral CA1 networks across a range of network thresholds linearly increasing from 0 to 0.385 (for each threshold: *p* < 0.005, two-sided Wilcoxon rank-sum test, ndCA1 = 335 units, nvCA1 = 228 units). Error bars denote standard error of the mean.

Although we found no significant differences in the mean pairwise correlations for dorsal and ventral CA1 (Figures 3C and 3D), these averages across all pairs of cells masked the differences in the structure of correlations, as evidenced by the increased variance in dorsal CA1 correlations (Figure 3C). To overcome this limitation, we visualized the structure of interactions by building network graphs, in which nodes denoted individual neurons and edges denoted a strong correlation between a pair of neurons (Figures 3E and 3F). To understand how correlations were distributed across the population, we calculated the degree, or number of incident edges, for each node (Figure 3G). Nodes in dorsal CA1 networks had significantly larger degrees than those in ventral CA1 networks, indicating that dorsal CA1 was more likely to contain highly connected “hub” neurons than ventral CA1. Importantly, regardless of the correlation threshold used to determine the presence of a network edge, the mean network degree of dorsal CA1 was larger than that of ventral CA1 (Figure 3H). Using this graph approach, we identified a novel difference in structure of correlations across neurons in the two regions, one that resulted in a less-connected network in ventral CA1.

### Decreased Dimensionality in Ventral CA1 Population Activity

The data collectively suggest that essential features of the hippocampal representation of space are encoded in the joint activity of neurons and that these activity patterns that cannot easily be summarized in metric such as place field size. To address this question, we explored the diversity of population activity states seen in our ensemble recordings. We defined a state of activity as any specific firing pattern across the population at a given time. These activity patterns varied as the animals moved through the virtual environment, thus changing the state of the network. We visualized these states (Figure 4A) using principle component analysis (PCA), a dimensional reduction method (Luczak et al., 2009; Yu et al., 2009) that also allowed us to calculate how covariations in the population activity shaped the dimensionality of the neuronal code. First, a three-dimensional projection of population activity (53 neurons in the dorsal CA1 example and 48 neurons in the ventral CA1 example) revealed complex trajectories corresponding to the evolution of activity as the animal traversed the track. Furthermore, in both the dorsal and ventral CA1 examples, the population activity occupied a different part of the space when the animal was running (indicated by the colored points) than when it was stationary (indicated by the grey points), consistent with the idea that the behavioral state of the animal shapes its neural activity (Chockanathan et al., 2020, 2021; Dadarlat and Stryker, 2017; Dipoppa et al., 2018; Niell and Stryker, 2010; Vinck et al., 2015).

**Figure 4.**
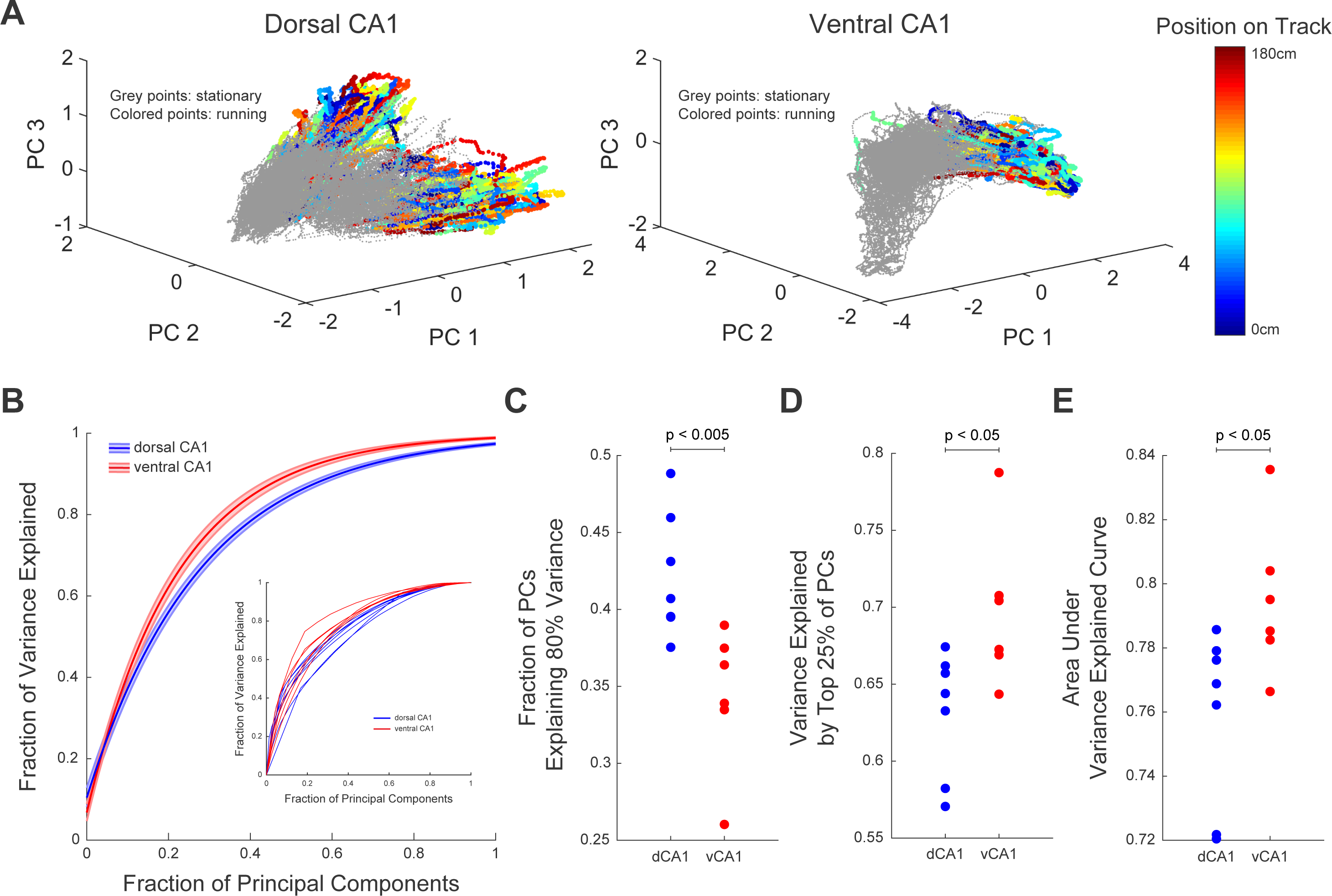
Dimensionality of Population Activity is Reduced in Ventral CA1. (A) Projections of population firing rates onto low-dimensional space defined by first three principal components for a representative dorsal CA1 (left) and ventral CA1 (right) recording session. (B) Cumulative explained variance as a function of fraction of total principal components ordered by their corresponding eigenvalues. Shaded regions denote standard error of the mean. Inset: Explained variance curves for single recording sessions. (C) A larger fraction of principal components was required to explain 80% of the variance in dorsal CA1 population than in ventral CA1 populations (mean ± std: dCA1 = 0.422 ± 0.040, vCA1 = 0.344 ± 0.046, *p* = 0.035, two-sided Wilcoxon rank-sum test, ndCA1 = 7 recording sessions, nvCA1 = 6 recording sessions). Each point denotes a recording session. (D) The first 25% of principal components explain a larger fraction of the variance in ventral CA1 than in dorsal CA1 (mean ± std: dCA1 = 0.632 ± 0.040, vCA1 = 0.697 ± 0.050, *p* = 0.035, two-sided Wilcoxon rank-sum test, ndCA1 = 7 recording sessions, nvCA1 = 6 recording sessions). Each point denotes a recording session. (E) The area under the variance explained curve was larger for ventral CA1 populations than for dorsal CA1 populations (mean ± std: dCA1 = 0.759 ± 0.027, vCA1 = 0.795 ± 0.024, *p* = 0.035, two-sided Wilcoxon rank-sum test, ndCA1 = 7 recording sessions, nvCA1 = 6 recording sessions). Each point denotes a recording session.

Although we observed complex trajectories of population activity in both dorsal and ventral CA1 in the same virtual environment (Figure 4B), the fraction of principal components required to explain 80% of the variance in the data was larger for dorsal than ventral CA1 (Figures 4C and S2). Additionally, the fraction of the total variance explained by the top 25% of principal components, as well as the area under the variance explained curve, was larger for ventral CA1 (Figures 4D and 4E). The population activity in ventral CA1 had fewer states, and therefore could be more easily summarized by a small number of variables than that of dorsal CA1. Taken together with previous results, this suggests that the ensemble activity of dorsal CA1 resides in a higher dimensional space than that of ventral CA1.

### Decreased Diversity of Population Activity Patterns in Ventral CA1

The results of Figures 3 and 4 indicate that the dimensionality of neural activity, and therefore the complexity of ensemble activity was larger in dorsal CA1 than ventral CA1. To understand what features of this activity conferred this increased complexity, we dissected the statistical properties of the population firing patterns in each region. To do this, we first defined a pattern of ensemble activity as a vector of neurons where an active cell was denoted with a 1 and an inactive cell denoted with a 0. In a given 10ms window then, a pattern or state of activity was defined as a vectors of 1s and 0s (Figure 5A) (Luczak et al., 2009; Miller et al., 2014; de Ruyter van Steveninck et al., 1997; Schneidman et al., 2006). We then compared the probability distributions of patterns from 18 neuron populations (made by subsampling from all recorded cells in a given recording session) in dorsal and ventral CA1 (Figures 5B and 5D). As illustrated by these representative examples, the pattern probability distribution was broader in dorsal CA1 than in ventral CA1, indicating a more diverse set of patterns. In addition, we found that the patterns that occurred in dorsal CA1 had more coactive neurons than those in ventral CA1 (Figures 5C, 5E, and 5F). The increased occurrence of such highly synchronous firing patterns allowed dorsal CA1 populations to occupy regions of the state landscape that were not observed with ventral CA1 populations, a potential mechanism contributing to the higher dimensionality of dorsal CA1 population activity (Figure 4).

**Figure 5.**
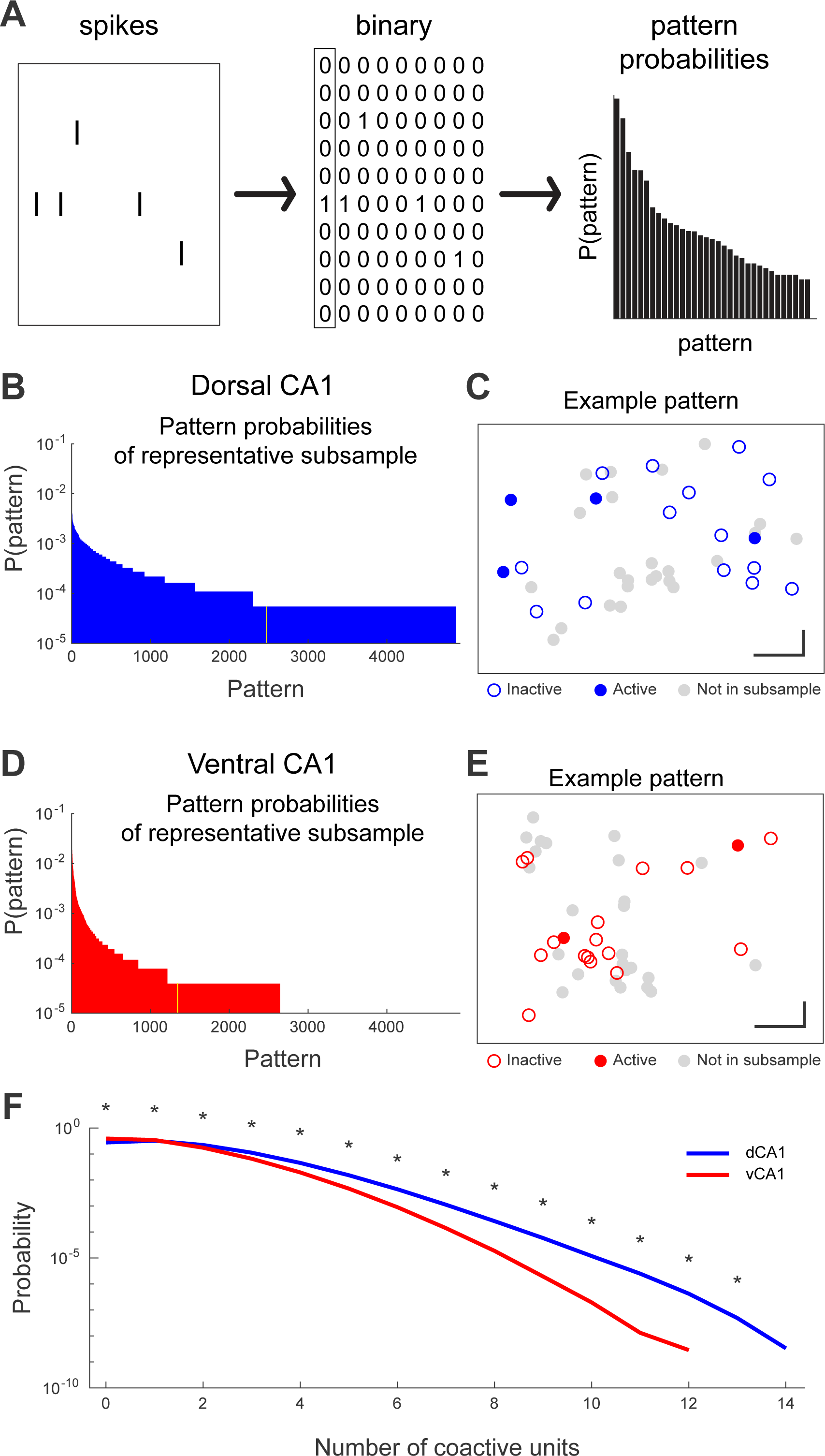
More Diverse and Complex Population Patterns in Dorsal CA1. (A) Schematic for calculating pattern probabilities. Left: the spike rasters for each unit were binned using 10ms non-overlapping windows and assigned a value of 0 if no spikes were present and a value of 1 if one or more spikes were present. Center: the combination of these binary states across all neurons in the subsample in a given window was denoted as a pattern. Right: Histogram of pattern probabilities. (B and D) Distribution of pattern probabilities for a single 18-unit subsample of a dorsal CA1 (B) and ventral CA1 (D) recording session. Yellow bar denotes the median probability pattern depicted in (C and E). (C and E) Visualization of a single representative pattern from dorsal CA1 (C) and ventral CA1 (E). Points denote the approximate physical position of individual units. Closed circles denote active units (1), open circles denote inactive units (0), and grey circles denote units not included in the current 18-unit subsample. (F) Mean probability of patterns grouped by number of coactive units, generated using 1000 18-unit subsamples in each recording session. Asterisks denote significant difference between dorsal and ventral CA1 probabilities (for each pattern type: *p* < 10^-6^, two-sided Wilcoxon rank-sum test, ndCA1 = 6000 samples, nvCA1 = 4000 samples).

### Population Activity in Ventral CA1 is Better Explained by Pairwise Interactions than in Dorsal CA1

The functional connectivity of neuronal populations, the number of states occupied by those populations, and the dimensionality of the neural code each reflect an underlying feature of CA1. By extension, properties such as functional connectivity, which reflect how neurons interact with one another, should also constrain the states that are occupied by the populations as well as the dimensionality of the neural code. We wished to know whether we could predict the frequency of activity patterns observed in groups of neurons by simply looking at the functional interactions between pairs of neurons. To what extent is the global structure of the network defined by these pairwise interactions?

One way of answering this question is the maximum entropy model, which explains the probability of all observed firing patterns of large populations of neurons with as little structure as possible (Figure 6A). If, for example, pairwise interactions were sufficient to explain the global structure of population activity, then a second order maximum entropy model would be able to predict population activity patterns based only on the firing rates of single neurons, reflected in the *hi* terms, and on the functional interactions between pairs of neurons, reflected in the *Jij* terms (Chockanathan et al., 2020; Meshulam et al., 2017; Schneidman et al., 2006; Shlens et al., 2006). When we compared the predicted and empirical distributions as a scatter plot for a representative 18-neuron subsample (Figures 6B and 6C), the maximum entropy model for the dorsal CA1 population showed many more points further away from the unity line than ventral CA1, indicating a higher error in prediction. Using the Kullback-Liebler divergence (KLD) a measure of the goodness of fit, we found that, across all pattern lengths we considered, the KLD was significantly larger for dorsal CA1 than ventral CA1 (Figures 6D and 6E). Pairwise interactions were better able to predict the state of the overall population in ventral CA1 than dorsal CA1, suggesting that pairwise correlations had significantly less explanatory power in dorsal CA1 as compared to ventral CA1. This was despite the fact that the values of the *hi* and *Jij* parameters were not significantly different for the two regions (Figure S3), consistent with our previous observation that neither the mean firing rates nor the mean pairwise correlations were significantly different for dorsal and ventral CA1 (Figures 2J and 3D). Thus, while Figures 4 and 5 show how the population activity in dorsal CA1 is more complex and diverse than that of ventral CA1, the results of our maximum entropy models demonstrate that this increased dimensionality cannot be explained simply by changes the structure of firing rates and pairwise interactions.

**Figure 6.**
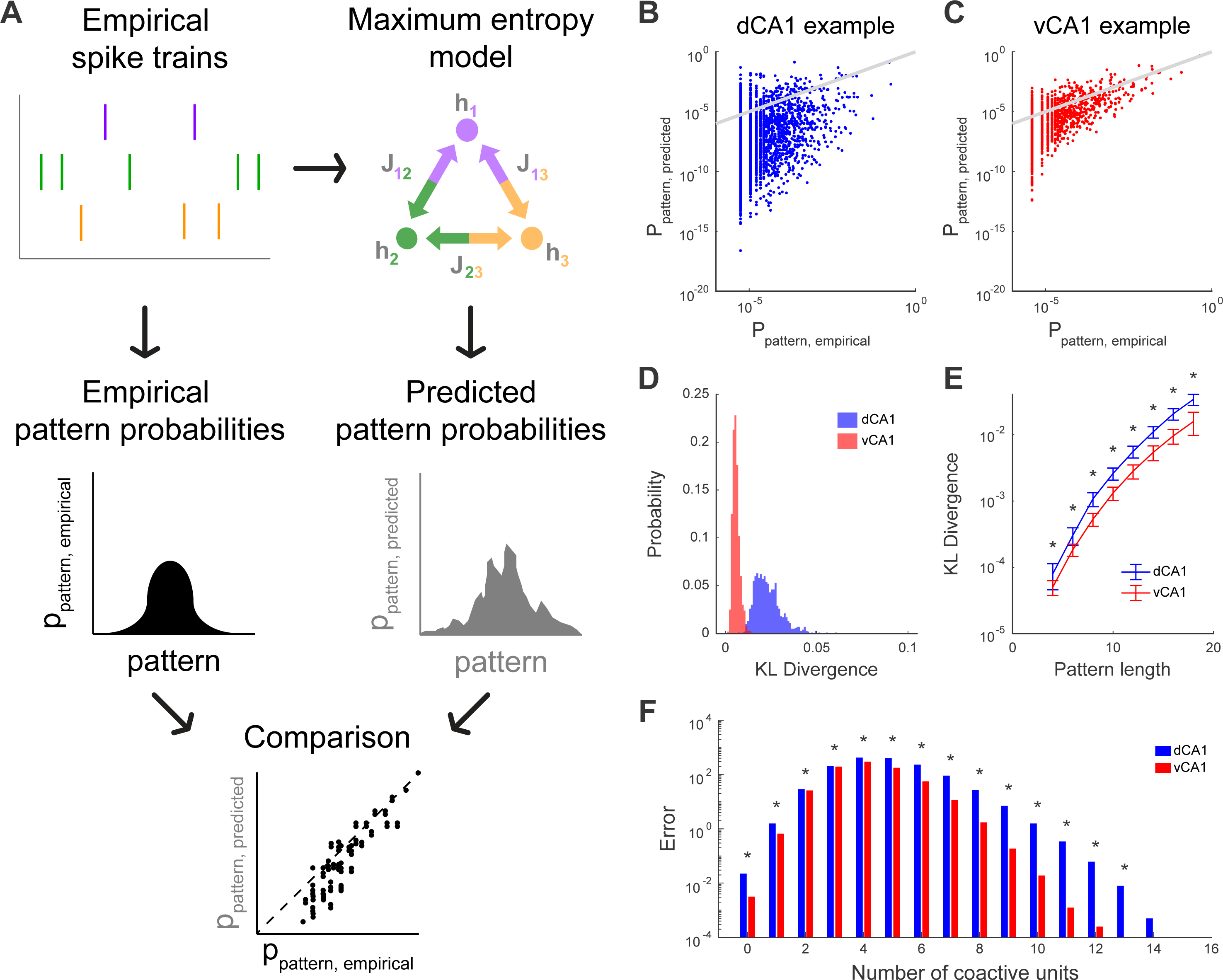
Pairwise Maximum Entropy Models Predict Population Patterns better in Ventral CA1 than in Dorsal CA1. (A) Schematic of maximum entropy models. Every neuron is assigned an activity term *hi* and every pair of neurons was assigned an interaction term *Jij*. Both terms were fit from the empirical spiking data. The models were then used to estimate the probability of every population pattern solely on the basis of these terms. The predicted pattern probabilities were then compared to the respective empirical probabilities. (B and C) Empirical pattern probabilities and maximum entropy model predicted pattern probabilities for a representative 18-unit subsample of a dorsal CA1 (B) and ventral CA1 (C) recording session. Each point denotes a single pattern and gray line denotes unity. (D) Distributions of Kullback-Liebler (KL) Divergence generated from 1000 random samples of 18-unit subpopulations in a representative dorsal and ventral CA1 recording session. (E) KL Divergence was larger for dorsal CA1 populations than ventral CA1 populations across a range of subpopulation sizes from 4 to 24 units. Asterisks denote significant difference between dorsal and ventral CA1 populations (for each pattern length: *p* < 10^-6^, two-sided Wilcoxon rank-sum test, ndCA1 = 100 samples per recording session, nvCA1 = 100 samples per recording session). Error bars denote standard error of the mean. (F) Pattern probability prediction error of maximum entropy model, grouped by number of coactive units, generated using 1000 18-unit subsamples in each recording session. Asterisks denote significant differences between dorsal and ventral CA1 prediction errors (for each pattern type: *p* < 10^-6^, two-sided Wilcoxon rank-sum test, ndCA1 = 6000 samples, nvCA1 = 4000 samples).

Instead, these data suggested that, in dorsal CA1, groups of neurons were co-active more often than would be predicted by their pairwise interactions. Indeed, when we then separated the patterns by the number of coactive units and analyzed the prediction error in each pattern class (Figure 6F), we found that the prediction errors in dorsal CA1 were larger than those in ventral CA1. Importantly, this gap was especially pronounced for patterns with high numbers of coactive units, suggesting that such highly synchronous patterns occurred far more frequently than would be predicted by pairwise functional coupling. Consistent with this idea, other groups have suggested that the coactivation of these groups of neurons may be orchestrated by higher order interactions (Ohiorhenuan et al., 2010). Our data suggest that these higher order interactions contribute significantly to shape ensemble activity in dorsal CA1, but are relatively weak or absent in ventral CA1. Such high order interactions may arise from the increased number of hubs we observed in the functional network structure, thereby driving the increased dimensionality of dorsal CA1 population codes of space.

### Decreased Population-level Spatial Information in Ventral CA1

Our data thus far reveal an underlying link between the functional organization of activity and the statistical implications for that activity in terms of the dimensionality of neuronal codes along the dorsal-ventral axis of CA1. We finally wished to determine what these differences in the statistics of population codes might mean for the representation of space throughout the hippocampus. Previous work has demonstrated that the activity of dorsal CA1 populations can be used to encode and decode the position of an animal in an environment (Wilson and McNaughton, 1993; Ziv et al., 2013). While ventral CA1 neurons have larger place fields (Jung et al., 1994), comparatively little is known about how larger place field size in individual neurons affects the activity of populations, the efficiency of this population activity on encoding information about an animals position, and what, if anything, these different strategies reveal about the functional heterogeneity of the hippocampus. To address this gap, we used an information theoretic approach (de Ruyter van Steveninck et al., 1997) to calculate the spatial information of population patterns and compare these quantities between dorsal and ventral CA1.

For a neural population to represent multiple spatial positions in a cognitive map, the activity patterns must be diverse as the animal navigates its environment. For the map to be reliable, patterns must correspond, reproducibly, to the position of the animal. Two components shape the spatial information content of populations: (1) the total entropy, which reflects the total number of population patterns observed over all regions of space and (2) the noise entropy, which reflects the population patterns observed in a specific region of space (see Method details). The difference between these two quantities is the information of the population, which reflects the degree to which the occurrence of a particular pattern can be used to infer the animal’s spatial position.

We generated pattern probability distributions conditioned on the animal’s position (Figure 7A) and quantified their diversity using entropy. A high entropy denoted a broad, flat probability distribution in which many patterns occurred with roughly equal probabilities while low entropy denoted a narrow distribution dominated by a small number of patterns.

**Figure 7.**
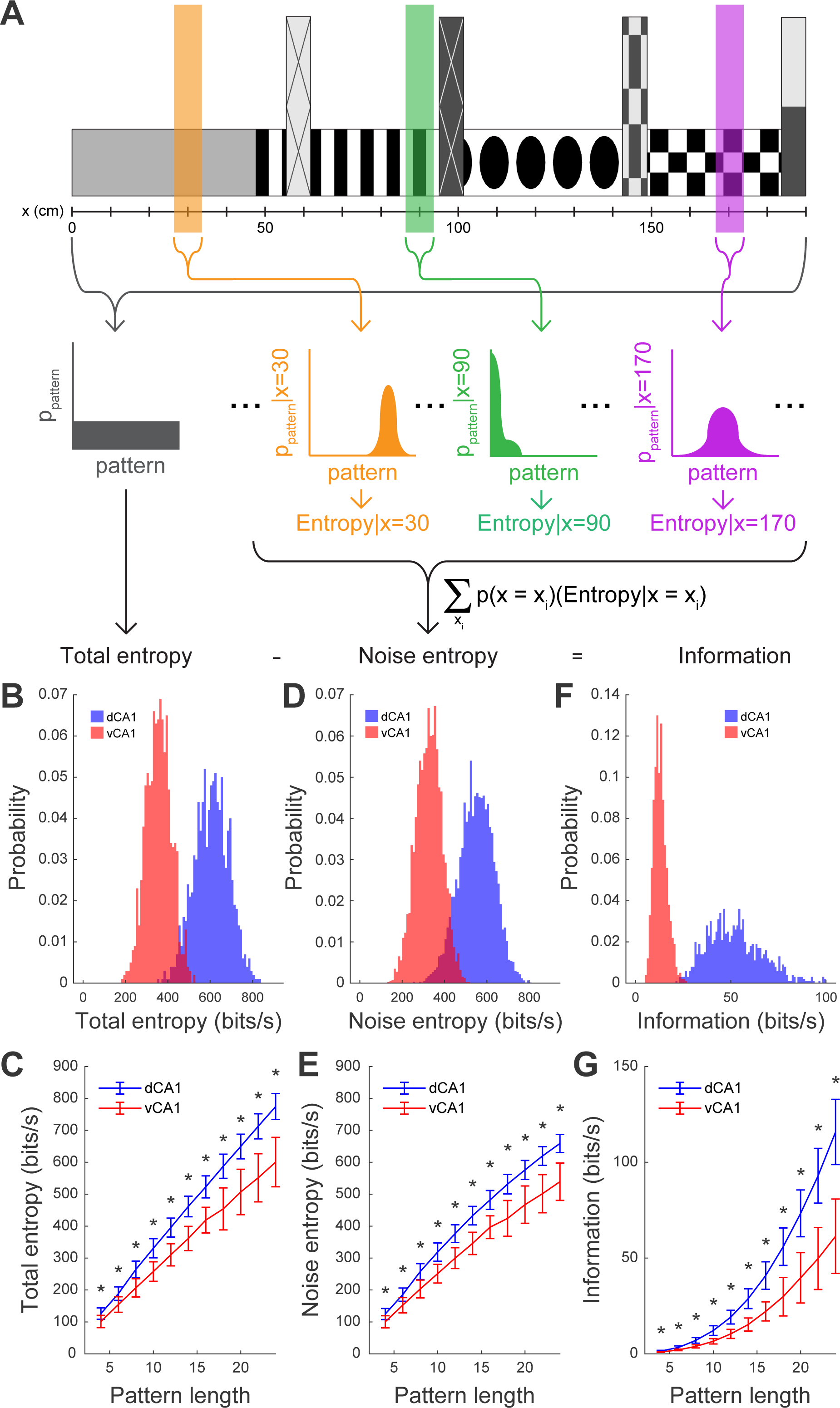
Dorsal CA1 Populations Collectively Encode more Spatial Information than Ventral CA1 Populations. (A) Schematic for calculating spatial information of CA1 populations. The conditional pattern probability distributions (yellow, green, and purple) were generated from the population patterns that were observed when the animal was at a particular position on the virtual track. The overall pattern probability distribution (grey) was generated by pooling the probabilities of all observed patterns, irrespective of the animal’s position. The entropy formula, ∑ −*p*(*log*_2_*p*) was then applied to each of the distributions. The total entropy was calculated from the overall distribution, while the noise entropy was calculated from conditional distributions, weighted by the amount of time the mouse spent at each position. The information was the difference between the total entropy and the information entropy. (B) Distributions of total entropy generated from 1000 random samples of 18-unit subpopulations in a representative dorsal and ventral CA1 recording session. (C) Total entropy was significantly larger for dorsal CA1 populations than ventral CA1 populations across a range of subpopulation sizes from 4 to 24 units. Asterisks denote significant difference between dorsal and ventral CA1 populations (for each pattern length: *p* <10^-6^, two-sided Wilcoxon rank-sum test, ndCA1 = 100 samples per recording session, nvCA1 = 100 samples per recording session). Error bars denote standard error of the mean. (D) Distributions of noise entropy generated from 1000 random samples of 18-unit subpopulations in a representative dorsal and ventral CA1 recording session. (E) Noise entropy was significantly larger for dorsal CA1 populations than ventral CA1 populations across a range of population sizes from 4 to 24 units. Asterisks denote significant difference between dorsal and ventral CA1 populations (for each pattern length: *p* < 10^-6^, two-sided Wilcoxon rank-sum test, ndCA1 = 100 samples per recording session, nvCA1 = 100 samples per recording session). Error bars denote standard error of the mean. (F) Distributions of spatial information generated from 1000 random samples of 18-unit subpopulations in a representative dorsal and ventral CA1 recording session. (G) Spatial information was significantly larger for dorsal CA1 populations than ventral CA1 populations across a range of population sizes from 4 to 24 units. Asterisks denote significant difference between dorsal and ventral CA1 populations (for each pattern length: *p* < 10^-6^, two-sided Wilcoxon rank-sum test, ndCA1 = 100 samples per recording session, nvCA1 = 100 samples per recording session). Error bars denote standard error of the mean.

We found that population spatial information was higher in dorsal CA1 than in ventral CA1 (Figures 7F and 7G), suggesting that the asymmetric representation of space found at the single unit level (Figure 2H) is preserved in populations. This increased information in dorsal CA1 was underpinned by a larger total entropy, suggesting that the overall number and diversity of observed patterns was larger in dorsal CA1 (Figures 7B and 7C), consistent with our previous results (Figure 5). Interestingly, however, we found that the noise entropy was also larger in dorsal than ventral CA1, suggesting that the diversity of patterns was larger in dorsal CA1 even within a single region of space (Figures 7D and 7E).

The findings of increased dimensionality shown in Figures 4 and 5 indicated that dorsal CA1 has a larger number of population patterns. The results in Figure 7 show that dorsal CA1 uses this increased neural “vocabulary” to represent space more effectively than ventral CA1. Taken together, our results provide additional evidence for the model of a functionally heterogeneous hippocampus, with dorsal CA1 specializing in spatial cognition as compared to ventral CA1.

## Discussion

We recorded the activity of neuronal populations in dorsal and ventral CA1 and examined the manner in which they collectively represented an animal’s position on a virtual track. Not only did we recapitulate earlier findings that individual neurons in ventral CA1 convey less information about position than those in dorsal CA1 (Jung et al., 1994),we also demonstrated that the collective activity of the population was differentially organized in the two regions. Dorsal CA1 populations formed more connected networks and showed more complex patterns of activity, which manifested as a larger total entropy, relative to ventral CA1 populations. This indication of increased dimensionality in dorsal CA1 activity was further supported by our finding that pairwise interactions were better able to predict population-level firing patterns in ventral CA1 than in dorsal CA1. Finally, by examining how the activity patterns in each region were organized by the position of the animal, we found that the population-level spatial information was also higher in dorsal CA1 than ventral CA1.

Along the dorsal-ventral axis of the hippocampus, there is a differentiation of functions that vary both in degree (for example, larger place fields in ventral CA1 than in dorsal CA1) as well as in kind (dorsal hippocampus is implicated in spatial memory, while ventral hippocampus is involved in social cognition and affect). These myriad cognitive and emotional roles are thought to arise from the variations in genetic (Cembrowski et al., 2016; Thompson et al., 2008), biophysical (Dougherty et al., 2012; Malik et al., 2016), circuit (Cenquizca and Swanson, 2007; Meira et al., 2018; Padmanabhan et al., 2019; Swanson and Cowan, 1977), and computational infrastructure (Ciocchi et al., 2015; Jimenez et al., 2018; Jung et al., 1994; Kjelstrup et al., 2008; Okuyama et al., 2016) across the dorsal-ventral axis of the hippocampus (Fanselow and Dong, 2010; Moser and Moser, 1998).

Our observation of decreased collective encoding of space in ventral CA1 mirrors broader topographic variations in spatial coding throughout the hippocampal formation. For grid cells, which have periodic spatial firing maps, the spacing and size of the grid vertices increase from the dorsal to the ventral boundaries of the medial entorhinal cortex (MEC) (Hafting et al., 2005; Stensola et al., 2012). A similar tuning gradient has been observed for head direction cells in layer III of the MEC, with dorsal neurons exhibiting sharper directional firing fields than ventral neurons (Giocomo et al., 2014). As part of the canonical trisynaptic loop, CA1 receives indirect inputs from entorhinal cortex as well as direct afferents from CA3. This circuit is organized in a lamellar fashion, such that dorsal or ventral entorhinal cortex preferentially projects to dorsal or ventral CA3, respectively, which in turn preferentially projects to dorsal or ventral CA1 (Andersen et al., 1971, 2000; Sloviter and Lømo, 2012). Our data suggest that, in addition to receiving differential inputs, the differences in functional interactions between neurons within dorsal and ventral CA1 may evidence the diverse ways in which spatial representations are encoded in the hippocampus.

This diversity may confer various computational benefits that support flexible behavioral strategies across diverse ethological environments. For instance, if place fields were fixed in size, then, as the territory an animal explores increased, more neurons would be required to comprehensively represent this expanding space. Indeed, previous studies in dorsal CA1 have demonstrated that as environments are expanded, more place cells are recruited and individual place cells show more numerous and enlarged place fields (Fenton et al., 2008). An increase in the size of the environment may require additional features such as hedonic value be represented in the hippocampus, so as to organize and structure an increasingly large environment. The larger place fields and coarser representations of ventral CA1 may be better suited for these parallel tasks. Studies of rats engaged in goal-directed spatial navigation tasks have found that place fields cover goal-related regions more densely than other regions (Fyhn et al., 2002; Hollup et al., 2001; Xu et al., 2018), while another study identified a dedicated population of neurons that are exclusively active when the animal visits a rewarded region. Perhaps the variation in spatial coding resolution between dorsal and ventral CA1 provides the flexibility to broadly map a large environment and simultaneously represent salient regions with very high precision. Future experiments that compare spatial representations in dorsal and ventral CA1 while animals explore multiple environments that vary in size and saliency would allow these hypotheses to be explicitly tested.

While we have focused on the hippocampal representation of space as an animal navigates a virtual environment, neurons in CA1 encode an array of non-spatial variables. In dorsal CA1, individual neurons have been identified that respond to time (MacDonald et al., 2011; Manns et al., 2007; Pastalkova et al., 2008), velocity (Czurkó et al., 2001; Góis and Tort, 2018), head-direction (Acharya et al., 2016; Leutgeb et al., 2000; Stefanini et al., 2020), and event sequences (Sun et al., 2020). Ventral CA1 neurons also exhibit diverse tuning properties for affective and social variables, including anxiety (Ciocchi et al., 2015; Jimenez et al., 2018) and conspecific identity (Deng et al., 2019; Okuyama et al., 2016; Rao et al., 2019). Often, neurons in both regions fire in response to combinations of multiple features (Ciocchi et al., 2015; Haimerl et al., 2019; Omer et al., 2018; Stefanini et al., 2020). Experimental and computational studies, both in hippocampus and primary sensory regions, such as the visual system and the main olfactory bulb, suggest that functional heterogeneity and mixed selectivity can increase the amount of information encoded by a population (Bernardi et al., 2020; Fusi et al., 2016; Padmanabhan and Urban, 2010; Shamir and Sompolinsky, 2006; Tripathy et al., 2013; Warland et al., 1997). Previous studies have demonstrated how this heterogeneity can improve the precision of spatial coding (Stefanini et al., 2020).

The diversity of population patterns we found in dorsal CA1 while the animal ran on the virtual track suggests that the activity of groups of neurons is marshalled in a way that is greater than the sum of their pairwise interacting parts to represent space. In parallel, in ventral CA1, the smaller number of patterns and lower spatial information may free up its coding “bandwidth” to represent other ethologically salient aspects of the world. We hypothesize that by differentially coordinating the activity of dorsal and ventral CA1 populations, the functional architecture of hippocampus supports distinct coding strategies, employing the coding space of dorsal CA1 to represent the animal’s position and leaving the capacity of ventral CA1 available for other emotive or affective behaviors.

## Acknowledgements

KP was supported by NSF CAREER (1749772), NIMH (R01MH11392), the Schmitt Foundation, and the Cystinosis Research Foundation. UC was funded by NIGMS (T32 GM007356).

## Author Contributions

K.P. supervised the project. K.P. and U.C. conceptualized the project, performed the analyses, made the figures, and wrote the manuscript. U.C. performed the experiments. Both authors approved the final version of the manuscript.

## STAR Methods

### KEY RESOURCES TABLE

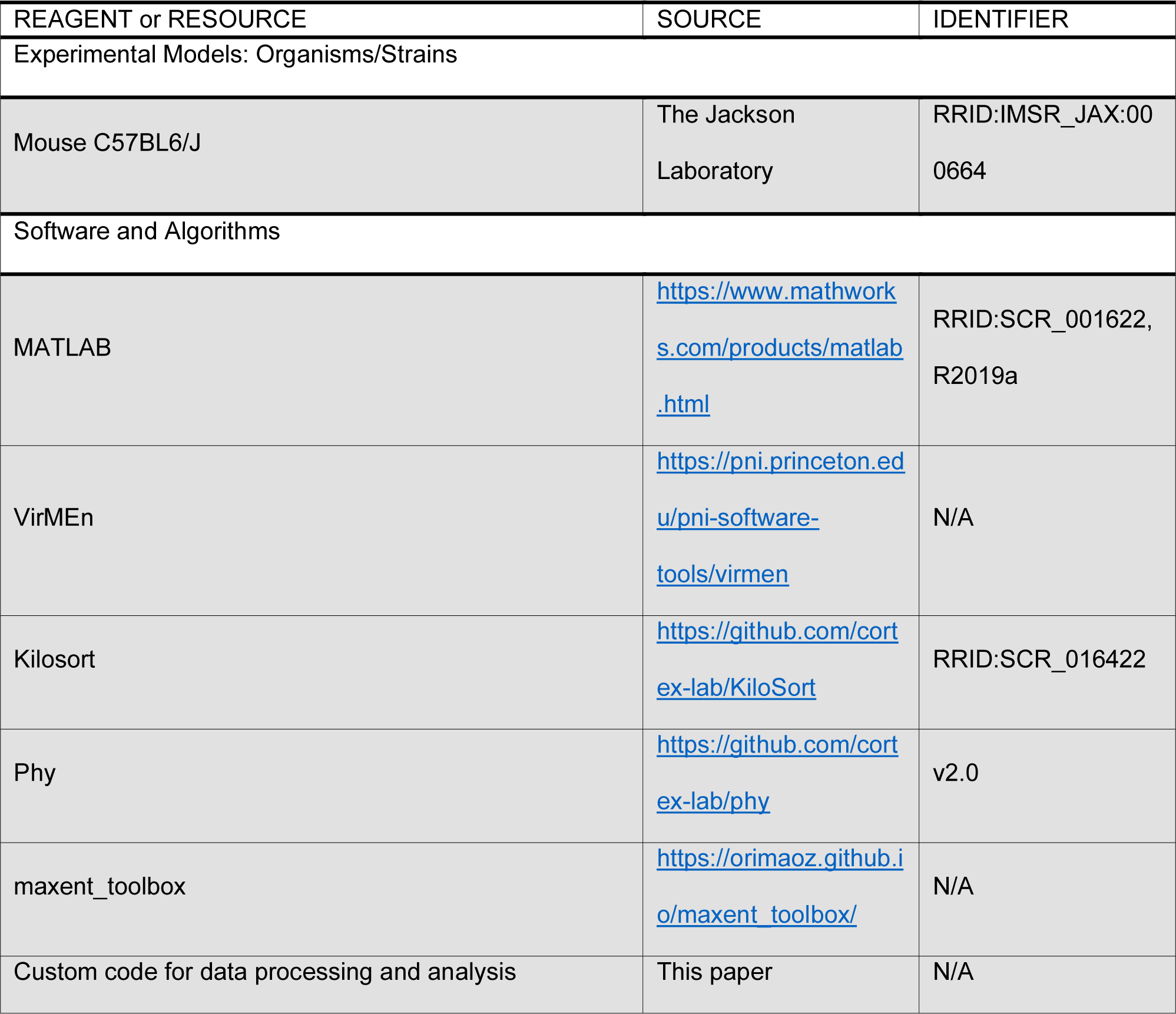

### RESOURCE AVAILABILITY

#### Lead Contact

Further information and requests for resources should be directed to the lead contact, Krishnan Padmanabhan.

#### Materials Availability

This study did not generate new reagents or animal lines.

#### Data and Code Availability

The datasets and code generated for this study are available upon reasonable request to the corresponding author.

### EXPERIMENTAL MODEL AND SUBJECT DETAILS

#### Mice

Four male C57BL6/J mice (The Jackson Laboratory, RRID:IMSR_JAX:000664) were included in this study. All mice were between 9 and 10 weeks of age at the time of recording. Mice were group-housed until headframe implantation, after which they were solo-housed. Mice were housed in transparent cages on a 12h:12h light/dark cycle. All recordings were performed in the light phase. Mice were healthy and were not used for any previous procedures. All procedures conformed to regulatory standards and were approved in advance by the Institutional Animal Care and Use Committee (IACUC) at the University of Rochester.

### METHOD DETAILS

#### Virtual Reality Setup

A virtual 1-dimensional track was created using the Virtual Reality MATLAB Engine (VirMEn) toolbox based on a previously published design (Gauthier and Tank, 2018; Meshulam et al., 2017). During the recording sessions, the position of the running wheel was transmitted to the computer by a 2-bit rotational encoder attached to the axle. This information was used to update the position of the animal on the virtual track, which was also saved. The image was projected onto a curved board and occupied approximately 180° of the visual field. When mice reached the end of the track, their virtual position was reset to the start of the track. The recording rig was enclosed in a box to minimize ambient light, odor, sound, and electromagnetic interference.

#### Head Fixing

Animals were anesthetized using a 1-2% isoflurane mixture and placed in a stereotactic surgical rig. The scalp was resected and the craniotomy sites for dorsal CA1 (coordinates relative to bregma: 2.5mm caudal, 1.5mm right) and ventral CA1 (coordinates relative to bregma: 3.15mm caudal, 3.15mm right).

Subsequently, a metal ground pin and custom 3D-printed headframe was affixed to the skull using dental cement (Ortho-Jet Powder and Jet Liquid, Lang Dental Mfg. Co., Wheeling, IL, USA) and veterinary adhesive (Vetbond, The 3M Company, Maplewood, MN, USA). Post-operative analgesia was provided for 72 hours with 0.03mL buprenorphine in accordance with approved protocols.

#### Run Training

During the 7-day period immediately following head frame implantation, animals were habituated to the virtual reality environment (Warner and Padmanabhan, 2020). Mice were head-fixed in the virtual environment and allowed to run for one hour each day. Run behavior was recorded during this training period, but electrophysiological recordings were not performed.

#### Electrophysiological Recording

Animals were anesthetized using a 1-2% isoflurane mixture and a craniotomy was performed over the right dorsal and ventral CA1 hippocampal subfields. Mice were then transferred to the running wheel and allowed to recover from anesthesia. A 128-channel nanofabricated electrode array (Du et al., 2011) was targeted to either dCA1 (coordinates: 2.5mm caudal, 1.5mm right, 1mm ventral) or vCA1 (coordinates: 3.15mm caudal, 3.15mm right, 4.25mm ventral). The virtual environment was activated and extracellular voltage recordings were collected at 30kHz in the 0.1-3500Hz frequency band. Recordings were performed for at least one hour at each recording site (for some animals, recordings were done at multiple regions of interest within dCA1 or vCA1). Afterwards, the electrode was moved to the other hippocampal subfield and the process was repeated (in half of the animals, dCA1 recordings were performed first, while in the other half, vCA1 recordings were performed first). The running behavior of the mouse and the position in the virtual track was also recorded simultaneously.

#### Spike Sorting

The open source automated spike sorting toolbox Kilosort (Pachitariu et al., 2016) was used to preprocess the electrophysiology recordings and identify the spike times of single units. The raw electrophysiology signal was high passed at 500Hz, the median signal from all channels was subtracted from each channel, and correlated noise across channels was removed. Subsequently, a set of template waveforms and spike times was generated and updated in an iterative manner to reconstruct the original data set. The waveforms and spike times at the end of this optimization process constituted putative single units. These were than manually curated using Phy (Rossant et al., 2016). Units were preserved, eliminated, or merged on the basis of their mean waveforms across multiple channels, amplitudes and inter-spike interval distributions. Units without clear refractory periods were excluded.

#### Run Behavior Analysis

The 2-bit wheel position information, recorded at 30kHz, was converted to a velocity using a window of size 150ms with 10ms time steps. Intervals during which the running velocity was greater than 5 cm/s were used for subsequent calculations (firing rates, correlations, entropy, spatial information, maximum entropy modeling, etc.). Only recording sessions during which the animal completed at least one full run along the virtual track were included in the analyses.

#### Single-Cell Spatial Tuning

The vector of each animal’s position on the track was sorted into 111 bins, each of size ∼1.7cm. For each unit, the mean firing rate in each bin was calculated. The resulting raw firing rate map was smoothed using a 5 bin-width square wave and then normalized such that the maximum and minimum for each unit was 1 and 0, respectively. The spatial information of each unit was then calculated with the following formula (Skaggs et al., 1993):

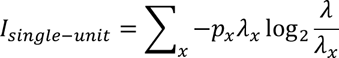

In this equation, *p*_*x*_ denotes the probability that the animal is occupying spatial bin *x*, λ_*x*_ denotes the firing rate in bin *x*, and λ denotes the mean firing rate across all bins. A complementary metric, sparsity, indicates the proportion of spatial bins over which a neuron is active (Skaggs et al., 1996):

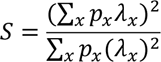

The center of a neuron’s place field was defined to the spatial bin in which the neuron’s firing rate was highest. The width of a place field was defined as the number of spatial bins around the place field center for which the neuron’s firing rate was at least one standard deviation higher than the mean.

### Firing Rate Correlations and Network Representations

A continuous firing rate trace of each neuron was generated by calculating the number of spikes in each 50ms interval of recording. The correlation coefficient between the firing rate traces of every pair of units in each animal was calculated. To convert the resulting correlation matrices to a graph, they were thresholded, such that the correlations that exceeded the threshold were preserved as an edge between the nodes representing that pair of neurons. The threshold was varied from 0 to 0.385 to cover the range of positive correlation strengths observed. For the network visualizations in Figures 3E and 3F (and for the pattern illustrations in Figures 5C and 5E), the physical location of the units was approximated by the location of the electrode contact on which the largest mean spike waveform of each unit was detected.

### Principal Components Analysis

The continuous firing rate traces used to calculate the correlations were also used to construct a covariance matrix. The eigenvectors of this matrix denoted the principal components, the axes along which the firing rates of the neuronal population showed the largest variance, and the eigenvalues denoted the variance along each principal component. For the scatter plots of low-dimensional population activity in Figures 4A and 4B, the firing rate traces were smoothed using a sliding Gaussian function with a standard deviation of 1s. This smoothing was done for the visualizations only and was not used for the data in Figures 4C-4E and S2.

Because the number of neurons, and thus, the number of principal components, varied between recording sessions, the explained variance plots could not be directly compared across recording sessions. Instead, the explained variance was plotted against the *fraction* of the total principal components in each recording. The area under this curve was calculated directly. To approximate the fraction of principal components that explained 80% of the variance, a nonlinear least-squares curve was generated for each plot using the trust-region-reflective algorithm (Coleman and Li, 1996).

### Population Entropy and Information

The population spike rasters were placed into 10ms bins and binarized such that the bin would be assigned a value of 0 if no spikes occurred in that 10ms interval and a value of 1 if one or more spikes occurred. The combination of 0’s and 1’s from multiple units in the population in a given time bin constituted a pattern (Chockanathan et al., 2020; de Ruyter van Steveninck et al., 1997; Schneidman et al., 2006). The number of units used in this subsample was varied from 4 units (in which case there were 2^4^ = 16 possible binary patterns) to 24 units (2^24^ = 16777216 possible patterns). The total entropy of the population was calculated from the equation, in which *p*_*k*_ denotes the probability of the *k*^th^ pattern:

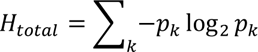

The total entropy describes the overall diversity of observed population patterns. However, the distribution of population patterns varied as a function of the animal’s position on the track. These conditional pattern probabilities (*p*_*k*_|*x*) were used to calculate the entropy as a function of animal’s position (the spatial bins used here were the same as those used to calculate the single-cell spatial information):

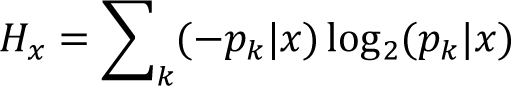

The average of all of these conditional entropies, weighted by the occupancy probability for each spatial bin, constitutes the noise entropy:

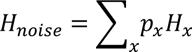

The spatial information of the population was calculated by the difference between the total entropy and the noise entropy:

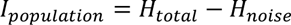

### Maximum Entropy Modeling

Maximum entropy models were fit to the data using the maxent_toolbox (Maoz and Schneidman, 2017). First, spike trains were binarized using 10ms non-overlapping bins, as in the entropy analysis. Second, subpopulations of n = 4 to 18 units were randomly generated. Third, for every neuron, a local field term *hi* was calculated. This term denotes the activity of that unit. Fourth, for every pair of units in the subsample, a pairwise interaction term *Jij* was calculated. This term describes the level of functional coupling between pairs of units. Subsequently, the probabilities of population patterns were predicted *using only these* hi *and* Jij *terms*:

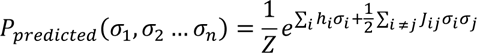

In this equation, σ_*i*_ indicates the binary state of the *i*^th^ unit and Z indicates the partition function used to normalize the probability distribution. Thus, the model attempts to predict the activity patterns of many-neuron populations without taking into account any higher-order interactions beyond pairwise couplings. The predicted pattern probabilities were then compared to the corresponding empirical probabilities and the prediction error was quantified using the Kullback-Liebler divergence (KLD).

### QUANTIFICATION AND STATISTICAL ANALYSIS

Statistical analyses were performed using MATLAB (MathWorks, Natick). For all hypothesis testing, two-sample Wilcoxon rank sum tests were performed, as the data were not assumed to be normally distributed. The alpha value was set at 0.05 and Bonferroni corrections were applied when multiple comparisons were performed. Details of statistical parameters (sample sizes used for statistical comparison, meaning of error bars, p-values) are reported in the figure legends or in the relevant section of the main text.

## Supplemental Information Titles and Legends

Figure S1. Association between Spatial Information, Firing Rate, and Place Field Location is preserved for Dorsal and Ventral CA1 Neurons

(A) Scatter plot of spatial information of each neuron against the location of its place field. There is no correlation between these variables in dorsal CA1 (*r* = -0.0044, *p* = 0.9366, Fisher transformation with two-sided Z-test, n = 335 neurons) nor in ventral CA1 (*r* = 0.0047, *p* = 0.9467, Fisher transformation with two-sided Z-test, n = 228 neurons). Moreover, this correlation coefficient is not significantly different between dorsal and ventral CA1 (*rdCA1* = -0.0044, *rvCA1* = 0.0047, *p* = 0.9124, Fisher transformation with two-sided Z-test, ndCA1 = 335 neurons, nvCA1 = 228 neurons). Each point denotes a single neuron, the black line denotes the least-squares regression, and the shaded area denotes the 95% confidence interval of the regression line. The text box shows *r*, the Pearson correlation coefficient, and the corresponding *p*-value.

(B) Scatter plot of spatial information of each neuron against its mean firing rate. These variables are correlated in both dorsal CA1 (*r* = -0.1482, *p* = 0.0066, Fisher transformation with two-sided Z-test, n = 335 neurons) and ventral CA1 (*r* = -0.1302, *p* = 0.0496, Fisher transformation with two-sided Z-test, n = 228 neurons). However, this correlation coefficient is not significantly different between dorsal and ventral CA1 (*rdCA1* = -0.1482, *rvCA1* = -0.1302, *p* = 0.8336, Fisher transformation with two-sided Z-test, ndCA1 = 335 neurons, nvCA1 = 228 neurons). Each point denotes a single neuron, the black line denotes the least-squares regression, and the shaded area denotes the 95% confidence interval of the regression line. The text box shows *r*, the Pearson correlation coefficient, and the corresponding *p*-value.

Figure S2. Explained Variance Curves and Corresponding Nonlinear Least Squares Fits

(A) Raw explained variance curves (black circles) and corresponding nonlinear least squares fits (blue lines; equations in inset) for each of the 7 recording sessions in dorsal CA1.

(B) Same as (A) for each of the 6 recording sessions in ventral CA1.

Figure S3. Maximum Entropy Parameters were Similar for Dorsal and Ventral CA1

(A) Mean maximum entropy activity terms *hi* from 1000 samples of 18-unit subpopulations. There was not a significance in *hi* between dorsal and ventral CA1 populations (mean ± std: dCA1 = -3.322 ± 0.203, vCA1 = -3.737 ± 0.549, *p* = 0.48, two-sided Wilcoxon rank-sum test, ndCA1 = 6 recording sessions, nvCA1 = 4 recording sessions). Each point denotes a recording session.

(B) Mean maximum entropy activity terms *hi* across a range of subpopulation sizes from 4 to 18 units. For each pattern length, 1000 subsamples were performed per recording session. Error bars denote standard error of the mean.

(C) Mean maximum entropy pairwise interaction terms *Jij* from 1000 samples of 18-unit subpopulations.

There was not a significance in *Jij* between dorsal and ventral CA1 populations (mean ± std: dCA1 = 0.075 ± 0.136, vCA1 = 0.044 ± 0.036, *p* = 0.48, two-sided Wilcoxon rank-sum test, ndCA1 = 6 recording sessions, nvCA1 = 4 recording sessions). Each point denotes a recording session.

(D) Mean maximum entropy pairwise interaction terms *Jij* across a range of subpopulation sizes from 4 to 18 units. For each pattern length, 1000 subsamples were performed per recording session. Error bars denote standard error of the mean.

## References

Acharya, L., Aghajan, Z.M., Vuong, C., Moore, J.J., and Mehta, M.R. (2016). Causal Influence of Visual Cues on Hippocampal Directional Selectivity. Cell 164, 197–207.

Andersen, P., Bliss, T.V.P., and Skrede, K.K. (1971). Lamellar organization of hippocampal excitatory pathways. Exp. Brain Res. 13, 222–238.

Andersen, P., Soleng, A.F., and Raastad, M. (2000). The hippocampal lamella hypothesis revisited. Brain Res. 886, 165–171.

Bernardi, S., Benna, M.K., Rigotti, M., Munuera, J., Fusi, S., and Salzman, C.D. (2020). The Geometry of Abstraction in the Hippocampus and Prefrontal Cortex. Cell 1–14.

Cembrowski, M.S., Bachman, J.L., Wang, L., Sugino, K., Shields, B.C., and Spruston, N. (2016). Spatial Gene-Expression Gradients Underlie Prominent Heterogeneity of CA1 Pyramidal Neurons. Neuron 89, 351–368.

Cenquizca, L.A., and Swanson, L.W. (2007). Spatial organization of direct hippocampal field CA1 axonal projections to the rest of the cerebral cortex. Brain Res. Rev. 56, 1–26.

Chockanathan, U., Warner, E.J., Turpin, L., O’Banion, M.K., and Padmanabhan, K. (2020). Altered dorsal CA1 neuronal population coding in the APP/PS1 mouse model of Alzheimer’s disease. Sci. Rep. 10, 1– 12.

Chockanathan, U., Crosier, E.J.W., Waddle, S., Lyman, E., Gerkin, R.C., and Padmanabhan, K. (2021). Changes in pair-wise correlations during running reshapes global network state in the main olfactory bulb. J. Neurophysiol. in press.

Ciocchi, S., Passecker, J., Malagon-Vina, H., Mikus, N., and Klausberger, T. (2015). Selective information routing by ventral hippocampal CA1 projection neurons. Science 348, 560–563.

Coleman, T.F., and Li, Y. (1996). An interior trust region approach for nonlinear minimization subject to bounds. SIAM J. Optim. 6, 418–445.

Czurkó, A., Hirase, H., Csicsvari, J., and Buzsáki, G. (2001). Sustained activation of hippocampal pyramidal cells by ‘space clamping’ in a running wheel. Eur. J. Neurosci. 11, 344–352.

Dadarlat, M.C., and Stryker, M.P. (2017). Locomotion Enhances Neural Encoding of Visual Stimuli in Mouse V1. J. Neurosci. 37, 3764–3775.

Deng, X., Gu, L., Sui, N., Guo, J., and Liang, J. (2019). Parvalbumin interneuron in the ventral hippocampus functions as a discriminator in social memory. Proc. Natl. Acad. Sci. U. S. A. 116, 16583– 16592.

Dipoppa, M., Ranson, A., Krumin, M., Pachitariu, M., Carandini, M., and Harris, K.D. (2018). Vision and Locomotion Shape the Interactions between Neuron Types in Mouse Visual Cortex. Neuron 98, 602–615.e8.

Dougherty, K.A., Islam, T., and Johnston, D. (2012). Intrinsic excitability of CA1 pyramidal neurones from the rat dorsal and ventral hippocampus. J. Physiol. 590, 5707–5722.

Dougherty, K.A., Nicholson, D.A., Diaz, L., Buss, E.W., Neuman, K.M., Chetkovich, D.M., and Johnston, D. (2013). Differential expression of HCN subunits alters voltage-dependent gating of h-channels in CA1 pyramidal neurons from dorsal and ventral hippocampus. J. Neurophysiol. 109, 1940–1953.

Du, J., Blanche, T.J., Harrison, R.R., Lester, H.A., and Masmanidis, S.C. (2011). Multiplexed, High Density Electrophysiology with Nanofabricated Neural Probes. PLoS One 6, e26204.

Fanselow, M.S., and Dong, H.W. (2010). Are the Dorsal and Ventral Hippocampus Functionally Distinct Structures? Neuron 65, 7–19.

Fenton, A.A., Kao, H.-Y., Neymotin, S.A., Olypher, A., Vayntrub, Y., Lytton, W.W., and Ludvig, N. (2008). Unmasking the CA1 Ensemble Place Code by Exposures to Small and Large Environments: More Place Cells and Multiple, Irregularly Arranged, and Expanded Place Fields in the Larger Space. J. Neurosci. 28, 11250–11262.

Fusi, S., Miller, E.K., and Rigotti, M. (2016). Why neurons mix: High dimensionality for higher cognition. Curr. Opin. Neurobiol. 37, 66–74.

Fyhn, M., Molden, S., Hollup, S., Moser, M.-B., and Moser, E.I. (2002). Hippocampal Neurons Responding to First-Time Dislocation of a Target Object. Neuron 35, 555–566.

Gauthier, J.L., and Tank, D.W. (2018). A Dedicated Population for Reward Coding in the Hippocampus. Neuron 99, 179–193.e7.

Giocomo, L.M., Stensola, T., Bonnevie, T., Van Cauter, T., Moser, M.-B., and Moser, E.I. (2014). Topography of Head Direction Cells in Medial Entorhinal Cortex. Curr. Biol. 24, 252–262.

Góis, Z.H.T.D., and Tort, A.B.L. (2018). Characterizing Speed Cells in the Rat Hippocampus. Cell Rep. 25, 1872–1884.e4.

Hafting, T., Fyhn, M., Molden, S., Moser, M.B., and Moser, E.I. (2005). Microstructure of a spatial map in the entorhinal cortex. Nature 436, 801–806.

Haimerl, C., Angulo-Garcia, D., Villette, V., Reichinnek, S., Torcini, A., Cossart, R., and Malvache, A. (2019). Internal representation of hippocampal neuronal population spans a time-distance continuum. Proc. Natl. Acad. Sci. U. S. A. 116, 7477–7482.

Henke, P.G. (1990). Hippocampal pathway to the amygdala and stress ulcer development. Brain Res. Bull. 25, 691–695.

Hollup, S.A., Molden, S., Donnett, J.G., Moser, M.B., and Moser, E.I. (2001). Accumulation of hippocampal place fields at the goal location in an annular watermaze task. J. Neurosci. 21, 1635–1644.

Jimenez, J.C., Su, K., Goldberg, A.R., Luna, V.M., Biane, J.S., Ordek, G., Zhou, P., Ong, S.K., Wright, M.A., Zweifel, L., et al. (2018). Anxiety Cells in a Hippocampal-Hypothalamic Circuit. Neuron 97, 670– 683.

Jung, M.W., Wiener, S.I., and McNaughton, B.L. (1994). Comparison of Spatial Firing Characteristics of Units in Dorsal and Ventral Hippocampus of the Rat. J. Neurosci. 74, 7347–7356.

Keinath, A.T., Wang, M.E., Wann, E.G., Yuan, R.K., Dudman, J.T., and Muzzio, I.A. (2014). Precise spatial coding is preserved along the longitudinal hippocampal axis. Hippocampus 24, 1533–1548.

Kjelstrup, K.B., Solstad, T., Brun, V.H., Hafting, T., Leutgeb, S., Witter, M.P., Moser, E.I., and Moser, M.-B. (2008). Finite Scale of Spatial Representation in the Hippocampus. Science (80-.). 321, 140 LP – 143.

Kjelstrup, K.G., Tuvnes, F.A., Steffenach, H.-A., Murison, R., Moser, E.I., and Moser, M.-B. (2002). Reduced fear expression after lesions of the ventral hippocampus. Proc. Natl. Acad. Sci. 99, 10825– 10830.

Klüver, H., and Bucy, P.C. (1937). “Psychic blindness” and other symptoms following bilateral temporal lobectomy in Rhesus monkeys. Am. J. Physiol.

Leutgeb, S., Ragozzino, K.E., and Mizumori, S.J.Y. (2000). Convergence of head direction and place information in the CA1 region of hippocampus. Neuroscience 100, 11–19.

Luczak, A., Bartho, P., and Harris, K.D. (2009). Spontaneous Events Outline the Realm of Possible Sensory Responses in Neocortical Populations. Neuron 62, 413–425.

MacDonald, C.J., Lepage, K.Q., Eden, U.T., and Eichenbaum, H. (2011). Hippocampal “Time Cells” Bridge the Gap in Memory for Discontiguous Events. Neuron 71, 737–749.

Malik, R., Dougherty, K.A., Parikh, K., Byrne, C., and Johnston, D. (2016). Mapping the electrophysiological and morphological properties of CA1 pyramidal neurons along the longitudinal hippocampal axis. Hippocampus 26, 341–361.

Manns, J.R., Howard, M.W., and Eichenbaum, H. (2007). Gradual Changes in Hippocampal Activity Support Remembering the Order of Events. Neuron 56, 530–540.

Maoz, O., and Schneidman, E. (2017). maxent_toolbox: Maximum entropy toolbox for MATLAB, version 1.0.2.

Meira, T., Leroy, F., Buss, E.W., Oliva, A., Park, J., and Siegelbaum, S.A. (2018). A hippocampal circuit linking dorsal CA2 to ventral CA1 critical for social memory dynamics. Nat. Commun. 9, 1–14.

Meshulam, L., Gauthier, J.L., Brody, C.D., Tank, D.W., and Bialek, W. (2017). Collective Behavior of Place and Non-place Neurons in the Hippocampal Network. Neuron 96, 1178–1191.e4.

Miller, J.K., Ayzenshtat, I., Carrillo-Reid, L., and Yuste, R. (2014). Visual stimuli recruit intrinsically generated cortical ensembles. Proc. Natl. Acad. Sci. 111, E4053–E4061.

Moser, M., and Moser, E.I. (1998). Functional Differentiation in the Hippocampus. Hippocampus 619, 608–619.

Moser, M.B., Moser, E.I., Forrest, E., Andersen, P., and Morris, R.G. (1995). Spatial learning with a minislab in the dorsal hippocampus. Proc. Natl. Acad. Sci. 92, 9697–9701.

Niell, C.M., and Stryker, M.P. (2010). Modulation of Visual Responses by Behavioral State in Mouse Visual Cortex. Neuron 65, 472–479.

O’Keefe, J., and Dostrovsky, J. (1971). The hippocampus as a spatial map. Preliminary evidence from unit activity in the freely-moving rat. Brain Res. 34, 171–175.

Ohiorhenuan, I.E., Mechler, F., Purpura, K.P., Schmid, A.M., Hu, Q., and Victor, J.D. (2010). Sparse coding and high-order correlations in fine-scale cortical networks. Nature 466, 617–621.

Okuyama, T., Kitamura, T., Roy, D.S., Itohara, S., and Tonegawa, S. (2016). Ventral CA1 neurons store social memory. Science 353.

Omer, D.B., Maimon, S.R., Las, L., and Ulanovsky, N. (2018). Social place-cells in the bat hippocampus. Science (80-.). 359, 218–224.

Pachitariu, M., Steinmetz, N., Kadir, S., Carandini, M., and Harris, K.D. (2016). Kilosort: realtime spike-sorting for extracellular electrophysiology with hundreds of channels. BioRxiv 61481.

Padmanabhan, K., and Urban, N.N. (2010). Intrinsic biophysical diversity decorrelates neuronal firing while increasing information content. Nat. Neurosci. 13, 1276–1282.

Padmanabhan, K., Osakada, F., Tarabrina, A., Kizer, E., Callaway, E.M., Gage, F.H., and Sejnowski, T.J. (2019). Centrifugal Inputs to the Main Olfactory Bulb Revealed Through Whole Brain Circuit-Mapping. Front. Neuroanat. 12, 115.

Pastalkova, E., Itskov, V., Amarasingham, A., and Buzsáki, G. (2008). Internally Generated Cell Assembly Sequences in the Rat Hippocampus. Science (80-.). 321, 1322 LP – 1327.

Rao, R.P., von Heimendahl, M., Bahr, V., and Brecht, M. (2019). Neuronal Responses to Conspecifics in the Ventral CA1. Cell Rep. 27, 3460–3472.e3.

Rich, P.D., Liaw, H.-P., and Lee, A.K. (2014). Large environments reveal the statistical structure governing hippocampal representations. Science 814, 814–817.

Rossant, C., Kadir, S.N., Goodman, D.F.M., Schulman, J., Hunter, M.L.D., Saleem, A.B., Grosmark, A., Belluscio, M., Denfield, G.H., Ecker, A.S., et al. (2016). Spike sorting for large, dense electrode arrays. Nat. Neurosci. 19, 634–641.

de Ruyter van Steveninck, R.R., Lewen, G.D., Strong, S.P., Koberle, R., Bialek, W., Steveninck, R.R.D.R. Van, Lewen, G.D., Strong, S.P., Koberle, R., and Bialek, W. (1997). Reproducibility and Variability in Neural Spike Trains. Science 275, 1805–1808.

Schneidman, E., Berry, M.J., Segev, R., and Bialek, W. (2006). Weak pairwise correlations imply strongly correlated network states in a neural population. Nature 440, 1007–1012.

Scoville, W.B., and Milner, B. (1957). Loss of recent memory after bilateral hippocampal lesions. J. Neurol. Neurosurg. Psychiatry 20, 11–21.

Shamir, M., and Sompolinsky, H. (2006). Implications of Neuronal Diversity on Population Coding. Neural Comput. 18, 1951–1986.

Shlens, J., Field, G.D., Gauthier, J.L., Grivich, M.I., Petrusca, D., Sher, A., Litke, A.M., and Chichilnisky, E.J. (2006). The Structure of Multi-Neuron Firing Patterns in Primate Retina. J. Neurosci. 26, 8254–8266.

Skaggs, W.E., McNaughton, B.L., and Gothard, K.M. (1993). An Information-Theoretic Approach to Deciphering the Hippocampal Code. Adv. Neural Inf. Process. Syst. 1030–1037.

Skaggs, W.E., McNaughton, B.L., Wilson, M.A., and Barnes, C.A. (1996). Theta phase precession in hippocampal neuronal populations and the compression of temporal sequences. Hippocampus 6, 149– 172.

Sloviter, R., and Lømo, T. (2012). Updating the Lamellar Hypothesis of Hippocampal Organization . Front. Neural Circuits 6, 102.

Stefanini, F., Kushnir, L., Jimenez, J.C., Jennings, J.H., Woods, N.I., Stuber, G.D., Kheirbek, M.A., Hen, R., and Fusi, S. (2020). A Distributed Neural Code in the Dentate Gyrus and in CA1. Neuron 107, 703–716.e4.

Stensola, H., Stensola, T., Solstad, T., Frøland, K., Moser, M.-B., and Moser, E.I. (2012). The entorhinal grid map is discretized. Nature 492, 72–78.

Sun, C., Yang, W., Martin, J., and Tonegawa, S. (2020). Hippocampal neurons represent events as transferable units of experience. Nat. Neurosci.

Swanson, L.W., and Cowan, W.M. (1977). An autoradiographic study of the organization of the efferent connections of the hippocampal formation in the rat. J. Comp. Neurol. 172, 49–84.

Thompson, C.L., Pathak, S.D., Jeromin, A., Ng, L.L., MacPherson, C.R., Mortrud, M.T., Cusick, A., Riley, Z.L., Sunkin, S.M., Bernard, A., et al. (2008). Genomic Anatomy of the Hippocampus. Neuron 60, 1010– 1021.

Tripathy, S.J., Padmanabhan, K., Gerkin, R.C., and Urban, N.N. (2013). Intermediate intrinsic diversity enhances neural population coding. Proc. Natl. Acad. Sci.

Vinck, M., Batista-Brito, R., Knoblich, U., and Cardin, J.A. (2015). Arousal and Locomotion Make Distinct Contributions to Cortical Activity Patterns and Visual Encoding. Neuron 86, 740–754.

Warland, D.K., Reinagel, P., and Meister, M. (1997). Decoding visual information from a population of retinal ganglion cells. J. Neurophysiol. 78, 2336–2350.

Warner, E.J., and Padmanabhan, K. (2020). Sex differences in head-fixed voluntary running behavior in C57BL/6J mice. Eur. J. Neurosci. 51, 721–730.

Wilson, M.A., and McNaughton, B.L. (1993). Dynamics of the hippocampal ensemble code for space. Science 261, 1055–1058.

Xu, H., Baracskay, P., O’Neill, J., and Csicsvari, J. (2018). Assembly responses of hippocampal CA1 place cells predict learned behavior in goal-directed spatial tasks on the radial eight-arm maze. Neuron.

Yu, B.M., Cunningham, J.P., Santhanam, G., Ryu, S.I., Shenoy, K. V, and Sahani, M. (2009). Gaussian-Process Factor Analysis for Low-Dimensional Single-Trial Analysis of Neural Population Activity. J. Neurophysiol. 102, 614–635.

Ziv, Y., Burns, L.D., Cocker, E.D., Hamel, E.O., Ghosh, K.K., Kitch, L.J., Gamal, A. El, and Schnitzer, M.J. (2013). Long-term dynamics of CA1 hippocampal place codes. Nat. Neurosci. 16, 264.

